# Conflicts with diarrheal pathogens trigger rapid evolution of an intestinal signaling axis

**DOI:** 10.1101/2020.03.29.014761

**Authors:** Clayton M. Carey, Sarah E. Apple, Zoё A. Hilbert, Michael S. Kay, Nels C. Elde

**Affiliations:** Department of Human Genetics, University of Utah, Salt Lake City, UT, 84112, USA; Department of Biochemistry, University of Utah, Salt Lake City, UT, 84112, USA

## Abstract

The pathogenesis of infectious diarrheal diseases is largely attributed to enterotoxin proteins that disrupt intestinal water absorption, causing severe dehydration. Despite profound health consequences, the impacts of diarrhea-causing microbes on the evolutionary history of host species are largely unknown. We investigated patterns of genetic variation in mammalian Guanylate Cyclase-C (GC-C), an intestinal receptor frequently targeted by bacterial enterotoxins, to determine how hosts might adapt in response to diarrheal infections. Under normal conditions, GC-C interacts with endogenous guanylin peptides to promote water secretion in the intestine, but signaling can be hijacked by bacterially-encoded heat-stable enterotoxins (STa) during infection, which leads to overstimulation of GC-C and diarrhea. Phylogenetic analysis in mammals revealed evidence of recurrent positive selection in the GC-C ligand-binding domain in primates and bats, consistent with selective pressures to evade interactions with STa. Using *in vitro* assays and transgenic intestinal organoids to model STa-mediated diarrhea, we show that GC-C diversification in these lineages results in substantial variation in toxin susceptibility. In bats, we observe a unique pattern of compensatory coevolution in the endogenous GC-C ligand uroguanylin, reflecting intense bouts of positive selection at the receptor-ligand interface. These findings demonstrate control of water physiology as a previously unrecognized interface for genetic conflict and reveal diarrheal pathogens as a source of selective pressure among diverse mammals.

## Main Text

Many enteric pathogens enhance their growth and dissemination to new hosts by secreting enterotoxins that cause diarrhea^1^. Dehydration resulting from these host-enterotoxin interactions represents a significant cause of mortality and morbidity in human populations, particularly in children under the age of five^2^. The intestinal receptor Guanylate Cyclase-C (GC-C) is a frequent target of enterotoxins encoded by diverse bacterial pathogens. Under normal conditions, GC-C activity is stimulated by interactions with the endogenous peptides guanylin and uroguanylin, leading to an increase in intracellular cGMP levels in enterocytes lining the small intestine and colon^3^ (Figure 1A). Enterocyte cGMP levels regulate osmotic balance of the gut by promoting chloride secretion through cystic fibrosis transmembrane conductance regulator (CFTR) and inhibiting sodium import through sodium/hydrogen exchanger 3 (NHE3), causing water to flow into the intestinal lumen. During infection, GC-C signaling can be hijacked by bacterial pathogens that produce heat-stable enterotoxins (STa). These toxin peptides mimic sequence and structural features of guanylins to overstimulate cGMP production and cause severe watery diarrhea^4^, promoting rapid dissemination to new hosts. Given the central role of STa-GC-C interactions in the pathogenesis of many diarrheal infections, we hypothesized that GC-C might adapt resistance to STa variants as part of an ongoing evolutionary conflict with diarrheal bacteria.

**Figure 1:**
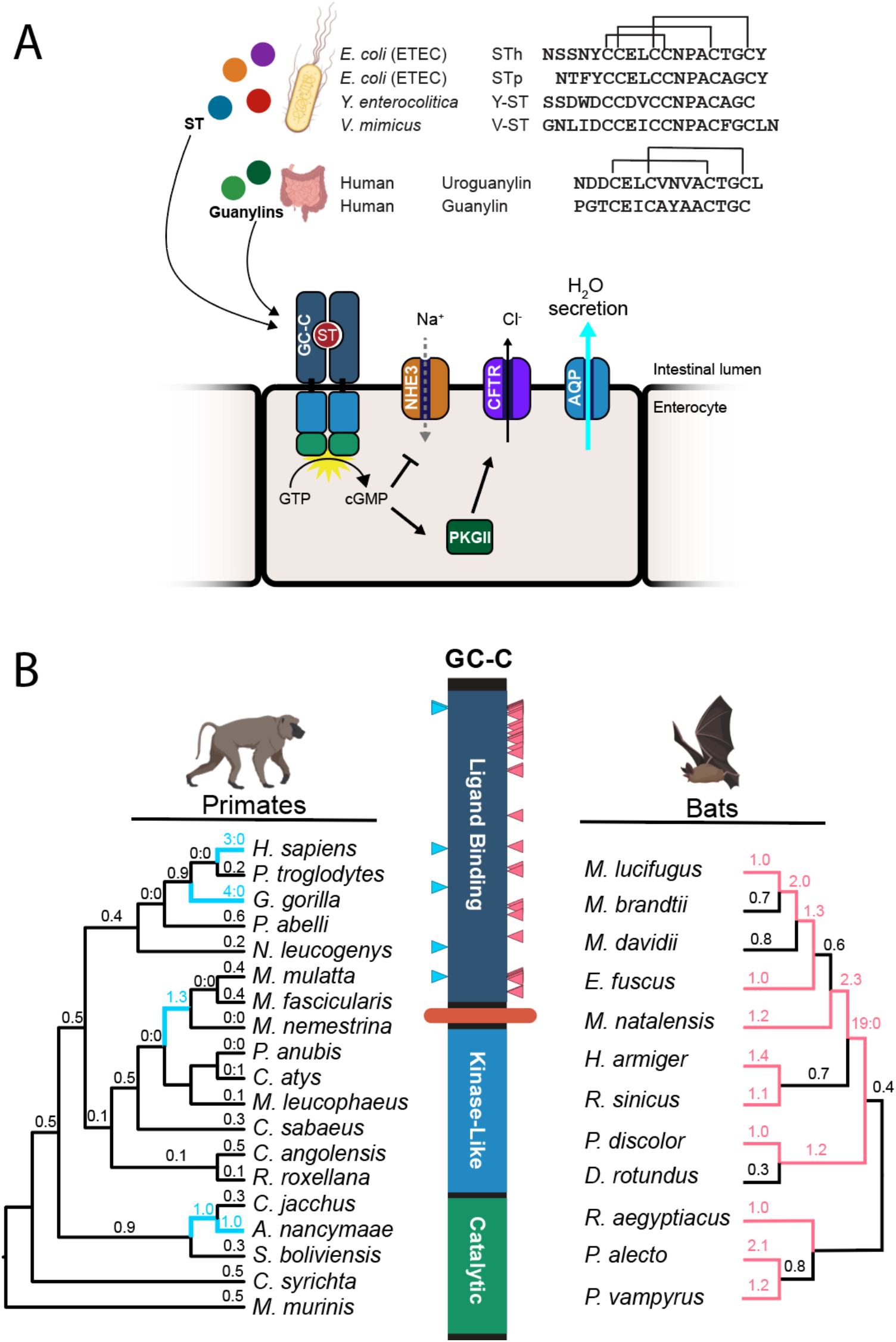
Rapid evolution of the heat-stable enterotoxin receptor in primates and bats. (A) Amino acid sequences of STa peptides encoded by four human pathogens are shown in comparison to human guanylin and uroguanylin. Disulfide bonds are indicated with connecting lines (top). Schematic of GC-C signaling in intestinal enterocytes. Activation of cGMP synthesis via ligand binding to GC-C results in chloride secretion through CFTR and inhibition of sodium transport via NHE3. Resulting shifts in ion balance facilitate water secretion into the intestinal lumen. Prolonged activation by STa causes excess water secretion and diarrhea (bottom). (B) Evolutionary analysis of GC-C in primates and bats. Phylogenetic trees indicate patterns of rapid evolution in primates (left) and bats (right). Branches with dN/dS >1 are highlighted in blue (primates) or magenta (bats) Codons with evidence of positive selection are indicated on the GC-C primary structure with blue (primates) or magenta (bats) triangles (PAML posterior probability >95%).

To determine whether GC-C evolved under selective pressure to modulate interactions with enterotoxins, we first collected the sequences of GC-C orthologs from 19 primate genomes for phylogenetic analysis (Table S1). Using statistical models that compare ratios of nonsynonymous to synonymous mutations (dN/dS) across species^5^,^6^, we find strong evidence of recurrent positive selection in GC-C (Table S2). Similar patterns of abundant nonsynonymous substitutions are widely reported for genes encoding dedicated immune functions that are targeted by pathogen factors^7^. This suggests that GC-C may similarly be subject to recurrent selection based on temporary advantages conferred by mitigating pathogen interactions. Unlike dedicated immune proteins, however, GC-C is functionally constrained by its physiological function of regulating water secretion through interactions with endogenous ligands. Indeed, the GC-C signaling axis is found in all vertebrates^8^, a pattern associated with functional conservation. To better understand the role of toxin interactions in GC-C evolution, we compared sequence evolution in the paralogous natriuretic peptide receptors (NPR1-2), which share similar overall structure and function with GC-C. NPR1 and NPR2 are not known to interact with pathogen-derived toxins^9^, and in contrast to GC-C, bear signatures of purifying selection with highly conserved sequences across primates (Tables S1, S2). Rapid evolution of GC-C is therefore associated with its unique interaction with pathogens.

Humans encounter a diverse array of STa peptides during infection with different bacterial pathogens. These commonly include toxin variants STp and STh encoded by strains of Enterotoxigenic *E. coli^10^*. Pathogenic strains from other bacterial genera, including *Yersinia* and *Vibrio* species, also encode distinct STa variants^11^,^12^. These peptides show considerable sequence variation in contrast to the endogenous GC-C ligands guanylin and uroguanylin that are nearly identical among primates (Figure 1A, S1). To further examine how GC-C evolution in primates might be influenced by toxin interactions, we mapped rapidly evolving codons with significantly elevated dN/dS ratios to the primary structure of GC-C. Strikingly, all six positively-selected sites mapped to the extracellular ligand-binding domain of GC-C, whereas the intracellular domain is highly conserved (Figure 1B, Table S3). These patterns of diversification of both STa and the GC-C ligandbinding domain support a model of recurring conflicts between diarrhea-causing bacteria and hosts over control of water secretion in the intestine.

In addition to primates, STa toxicity has been demonstrated in other mammalian species including rodents and livestock^4^. To test whether GC-C might be subject to recurrent positive selection in other mammalian lineages, we collected sequences of GC-C from four additional groups of mammals including *Bovidae* (cloven-hooved ruminants), *Caniformia* (suborder of dog-like carnivores), *Myomorpha* (mouse-like rodents), and *Chiroptera* (bats) (Table S1). In contrast to primates, substitution patterns among *Caniformia* and *Bovidae* are consistent with purifying selection, while signals of positive selection are supported in only a subset of tests among *Myomorpha* GC-C (Table S2).

Patterns of positive selection were remarkably widespread in bats and strongly supported by all phylogenetic tests. Examination of bat GC-C sequences revealed 35 codon positions with significantly elevated dN/dS values mapping exclusively to the GC-C ligand-binding domain (Figure 1B, Table S3). Sequence comparisons of bat GC-C and NPR2 ligand-binding domains revealed extensive diversification in GC-C relative to the closely related paralog (Figure S3). Although the structural basis of the interaction between GC-C and STa has not yet been determined, a putative ligand-binding pocket was proposed in a membrane-proximal region between residues 411–416 (ref ^13^). A cluster of seven sites with elevated dN/dS values among bats is in close proximity to this region, in addition to one site identified in our analysis of GC-C in primates (Figure S2).

Bats are known to harbor many types of viruses, including several human pathogens^14^. This widespread association with viruses is attributed to high density population structures and the ability of long-distance flight^15^. Notably, frequent infection with coronaviruses likely sparked high levels of sequence diversification in the viral entry receptor ACE-2 among bats^16^, illustrating the intensity of selection pathogens place on these species. In addition to viruses, bats can carry bacteria that act as human pathogens, including genera that encode STa^17^. Furthermore, STa variants have been detected in DNA isolated from the feces of wild bats^18^. Our detection of signals of intense positive selection in the GC-C ligand-binding domain of bats suggests that diarrheal pathogens may have profoundly impacted the evolution of bats, similar to their established coevolutionary relationship with viruses (Figure 1B).

To test if rapid evolution of GC-C ligand-binding domains result in functional differences in STa susceptibility, we generated cell lines stably expressing GC-C from seven primate and five bat species. Functional diversity in these GC-C variants was assessed by measuring intracellular cGMP production in response to chemically synthesized uroguanylin and STa variants from four strains of pathogenic bacteria. HEK293T cells expressing human, chimpanzee, gorilla, gibbon and African green monkey GC-C generated similar levels of cGMP upon stimulation with STa variants from pathogenic *E. coli* (STp and STh), *Yersinia enterocolitica* (Y-ST), and *Vibrio mimicus* (V-ST) (Figure 2A, S5). In contrast, cells expressing GC-C from orangutan and rhesus macaque were significantly more susceptible to STp, showing a large increase in cGMP production relative to other species (Figure 2A). Dose-response curves comparing maximal activation of human and orangutan GC-C to each toxin revealed similar maximal responses for human GC-C, while orangutan GC-C exhibited increases in relative vulnerability to STp and STh toxin variants compared to Y-ST and V-ST (Figure 2B, 2C). Thus, variation in GC-C among primates results in differing levels of susceptibility to diverse toxins, providing evidence that ancient STa-like peptides influenced the course of GC-C evolution.

**Figure 2:**
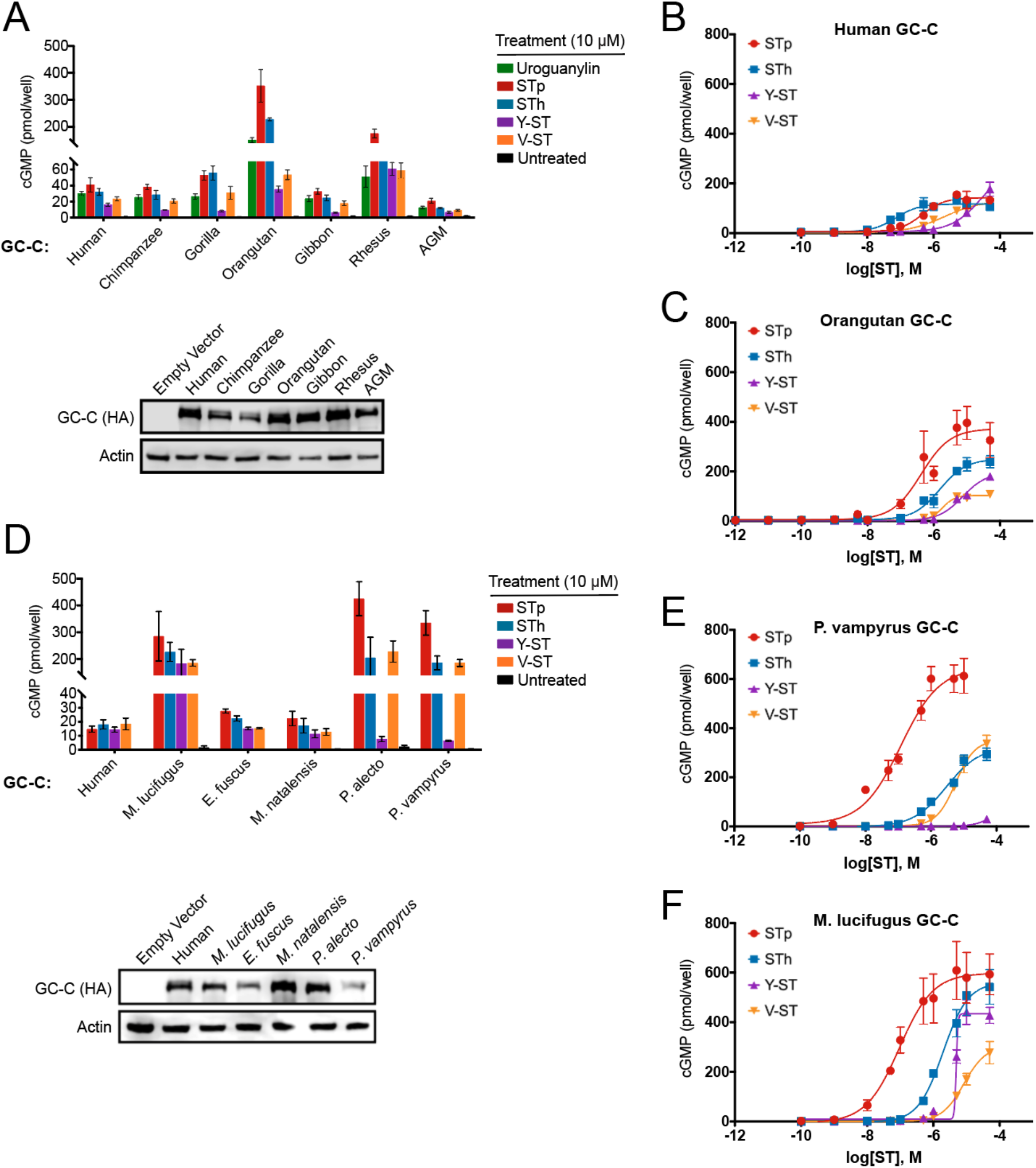
Species-specific patterns of STa susceptibility in primates and bats. (A and D) (Top) cGMP levels in HEK293T cells stably expressing GC-C from seven primate species (A) and five bat species (D) after treatment with 10 μM of the indicated peptide. (Bottom) Immunoblot of HA-tagged GC-C from indicated cell lines (B, C, E, F). Intracellular cGMP dose-response curves for the indicated ligands are shown in HEK293T cells expressing human (B), orangutan (C), *P. vampyrus* (E), and *M. lucifugus* (F) GC-C.

Differences in toxin susceptibility were even more pronounced among GC-C encoded by bats. We cloned GC-C from representatives of the vesper bats: *Myotis lucifugus* (little brown bat), *Eptesicus fuscus* (big brown bat), and *Miniopterus natalensis* (Natal long-fingered bat), as well as Old World fruit bats: *Pteropus vampyrus* (large flying fox) and *Pteropus alecto* (black flying fox) for expression in cell lines. This sampling captures roughly a 50 million year interval of divergence from the common ancestor of chiropteran bats^19^. We discovered a wide range of susceptibility to STa variants across bat species. While GC-C from the vesper bats *E. fuscus* and *M. natalensis* responded similarly to all four toxins at high concentrations and produced modest levels of cGMP comparable to human GC-C, cells expressing *M. lucifugus, P. vampyrus*, and *P. alecto* GC-C produced nearly 10-fold more cGMP in response to toxin treatments (Figure 2D-F). Intriguingly, both *P. vampyrus* and *P. alecto* failed to respond to treatment with 10 μM Y-ST in notable contrast to GC-C encoded by *M. lucifugus* (Figure 2D). Additional comparisons across a wide range of toxin concentrations revealed that *P. vampyrus* GC-C is only activated by Y-ST stimulation at concentrations exceeding 10 μM, well outside probable concentrations encountered during infection^20^ (Figure 2E). Thus, consistent with strong signatures of positive selection for GC-C from bats, susceptibility to STa variants widely differs between bat species, with some receptors appearing resistant to activation at physiologically plausible concentrations of toxin.

We next sought to more directly measure the physiological significance of differing toxin susceptibility we observed in our cGMP generation assays. Cultured organoids composed of intestinal enterocytes have recently emerged as a means to directly assay symptoms of secretory diarrhea^21^. In this system, GC-C mediated water secretion induces organoid swelling upon ligand stimulation, directly modeling symptoms of diarrhea. We cultured organoids derived from the small intestine of GC-C^-/-^ mice to investigate the relationship between cGMP production and water secretion across species. We chose to compare organoids expressing GC-C from humans and the fruit bat *P. vampyrus*, which displayed heightened susceptibility to STp, but marked resistance to Y-ST. While mouse GC-C^-/-^ organoids fail to respond to ligand stimulation, complementation with human or *P. vampyrus* GC-C via lentiviral transduction rescued swelling in response to STa (Figure 3A, 3B). Organoids expressing *P. vampyrus* GC-C displayed considerably increased water secretion during STp treatment as measured by the total change in organoid area compared to organoids expressing human GC-C (Figure 3A). Consistent with our *in vitro* analysis, treatment with Y-ST resulted in minimal swelling of organoids expressing *P. vampyrus* GC-C compared to human (Figure 3C, 3D). Thus, differences in susceptibility to STa variants likely have a direct influence on the level of water secretion and symptoms of diarrhea experienced by host species during infection.

**Figure 3:**
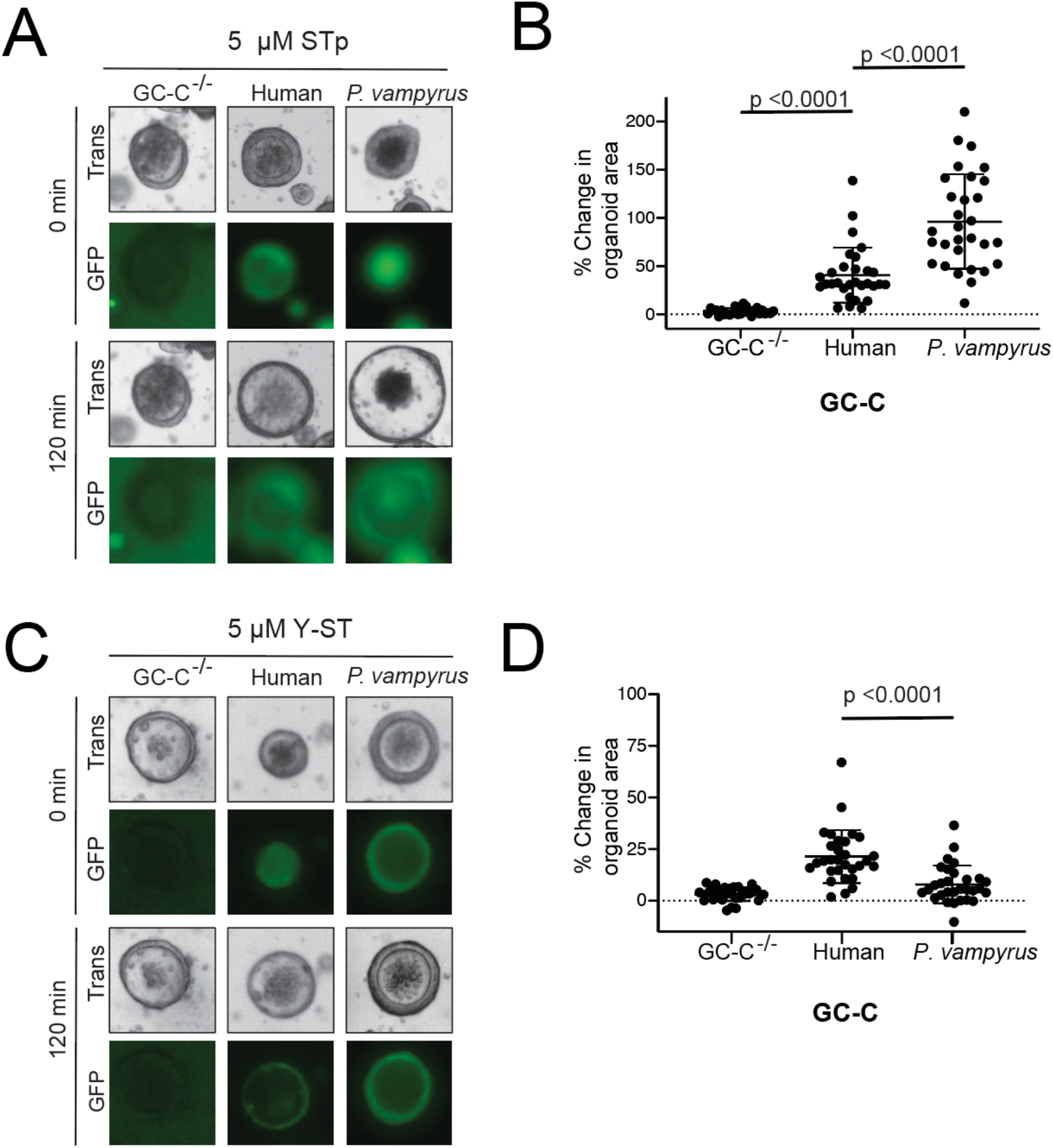
Intestinal organoids model physiology of toxin susceptibility. (A) Intestinal organoids were cultured from GC-C^-/-^ mice and transduced with lentivirus encoding GC-C variants and a GFP reporter. Representative images are shown of individual organoids expressing GC-C from the indicated species before and after 120 min of treatment with 5 μM STp. (B) Change in individual organoid area was quantified following 120 min of STp treatment for organoids expressing GC-C from the indicated species. (C) Representative images of individual organoids expressing GC-C from the indicated species before and after 120 min of treatment with 5 μM Y-ST. (D) Change in individual organoid area was quantified following 120 min of Y-ST treatment for organoids expressing GC-C from the indicated species.

In order to regulate intestinal water levels GC-C must interact with endogenous guanylin and uroguanylin peptides^8^. To determine how rapid divergence of GC-C might influence the evolution of its cognate ligands, we next examined diversity in uroguanylin, the more potent of the two peptides^22^. Sequence comparisons revealed that secreted uroguanylin peptides are generally highly conserved across mammals with little variation occurring outside the most N- and C-terminal residues (Figure S4). In bats, however, uroguanylin sequences are highly variable, with frequent mutations occurring in core residues of the peptide (Figure 4A). Given the variability in sequence and toxin interactions we observed in bat GC-C, we hypothesized that sequence variation in bat uroguanylin might reflect compensatory mutations required to maintain affinity for its rapidly evolving receptor.

**Figure 4:**
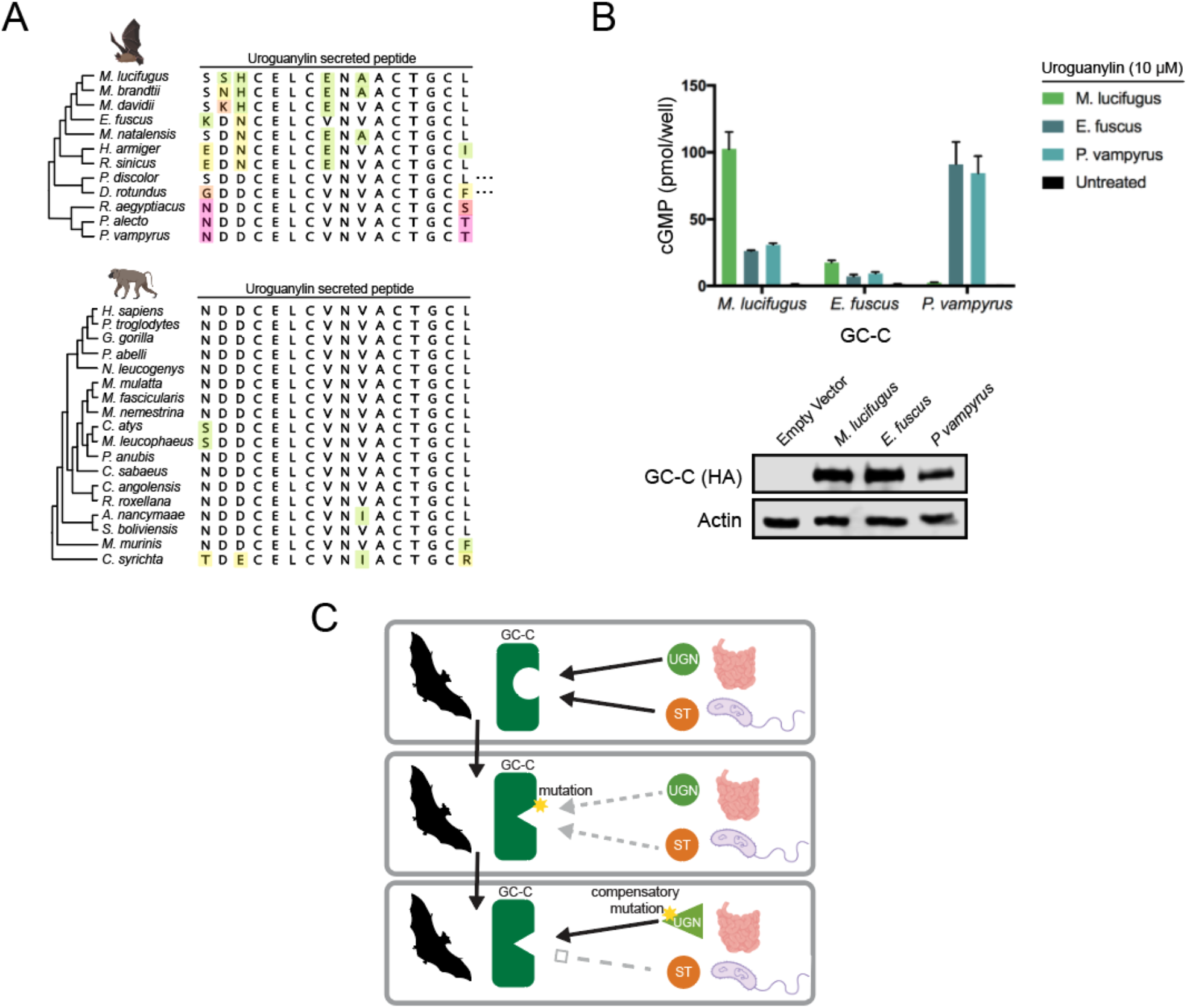
Coevolution of GC-C and uroguanylin in bats. (A) Amino acid alignment of the mature uroguanylin peptide based on genome sequences of the indicated bat (top) or primate (bottom) species. Residues with mutations relative to the consensus within each clade are uniquely colored. Elipses represent stop codon loss mutations in the indicated bat species. (B) (Top) cGMP levels in HEK293T cells expressing *P. vampyrus* or *M. lucifugus* GC-C after treatment with 10 μM uroguanylin from the indicated species. (Bottom) immunoblot of HA-tagged GC-C from the indicated cell lines. (C) Model for pathogen driven evolution of uroguanylin in bats. Mutations in GC-C that result in a loss of affinity for both STa and uroguanylin may provide a net fitness benefit, allowing time for compensatory mutations to arise.

To test if uroguanylin co-evolved with GC-C in bats, we synthesized active isomers of uroguanylin peptides encoded by the vesper bats *M. lucifugus* and *E. fuscus*, as well as the more distantly related *P. vampyrus* (Figure 4A, S6). Treatment of cells expressing *P. vampyrus* GC-C with uroguanylin from *M. lucifugus* did not stimulate cGMP production, whereas treatment with *P. vampyrus* uroguanylin robustly activates catalytic activity (Figure 4C). Conversely, *M. lucifugus* GC-C responds more strongly to its species-matched uroguanylin than to the *P. vampyrus* peptide (Figure 4B). Intriguingly, *E. fuscus* GC-C responds weakly to all uroguanylin variants tested, including its own peptide (Figure 4B). These experiments are consistent with compensatory co-evolution of GC-C and uroguanylin in bats, and also reveal that some species may exist in intermediate states with sub-optimal affinity for uroguanylin.

Given functionally consequential variation in GC-C that impacts both enterotoxin and uroguanylin interactions, we propose a model of compensatory coevolution triggered by bacterial enterotoxin interactions (Figure 4C). In this model, mutations in GC-C that allow escape from overstimulation by STa can provide a fitness benefit even at the cost of disrupting the interaction with uroguanylin. This intermediate state of low affinity with endogenous peptides is tolerated while toxin susceptible variants are culled from the population, given that survival is possible when GC-C signaling is disrupted^23^. Subsequent compensatory mutations in uroguanylin that optimize signaling interactions with GC-C might then outcompete mismatched variants with compromised receptorligand interfaces. In this proposed scenario, a single pathogen encoded protein directly influences the evolution of the host receptor, which subsequently impacts variation in the endogenous ligand.

Our work reveals how genes involved in intestinal water physiology can rapidly adapt in ongoing conflict with enteric pathogens. Similar to well established conflicts between host immune defenses and pathogen effectors, the recurrent evolutionary innovation seen in the GC-C ligand-binding domain indicates that modulating interactions with bacterial toxins has been critical for survival in primates and bats. We show that this diversification in the molecular machinery of the intestine likely contributes to disease susceptibility in host species and may restrict the host range of STa producing pathogens. Our observation of remarkable diversification at the GC-C receptor-ligand interface in bats further suggests that pathogens can spark compensatory coevolution within host signaling pathways. Together these findings illustrate the far-reaching impacts of deadly diarrheal infections on the evolutionary history of diverse mammals.

## Acknowledgements

The authors thank Scott Waldman and Amanda Pattison for generously providing GC-C knockout organoids. N.C.E. is supported by the National Institute of Health, United States (R01GM114514 and R35GM134936) and the Burroughs Wellcome Fund Investigators in the Pathogenesis of Infectious Disease Program, United States. M.S.K. is supported by the National Institute of Health, United States (P50AI150464). N.C.E and M.S.K. are H.A. & Edna Benning Presidential Endowed Chairs (University of Utah, United States). C.M.C is supported by the National Institute of Health, United States (T32AI055434).

## Author Contributions

C.M.C. and N.C.E. designed the study and wrote the manuscript. C.M.C and Z.A.H performed the evolutionary analysis. C.M.C. and Z.A.H. performed the cell culture and organoid experiments. S.E.A. and M.S.K. synthesized and purified all peptides. All authors reviewed and edited the manuscript.

## Methods

### Evolutionary analysis

GC-C and NPR1-3 nucleotide sequences were retrieved from NCBI GenBank for each species. Prior to analysis, sequences for each gene within each clade were first aligned using clustalW. In each analysis, a generally agreed upon phylogeny was used for each clade. Evidence for gene-wide positive selection was obtained using the branch-site models implemented in PAML and BUSTED software. In PAML, alignments were fitted to a F3×4 codon model and likelihood ratio tests were performed based on comparisons of NSsites model 2 to model 1 and model 8 to model 7. BUSTED analysis was performed on all branches of a user-specified tree using an online webserver with default parameters (https://www.datamonkey.org/busted). Gene trees based on the amino acid sequence of GC-C and NPR2 were generated by first aligning amino acid sequences with clustalW in each clade. Trees were generated using the PhyML plugin implemented in Geneious software using the Le Gascuel substitution model.

### Sequence alignment and figure generation

Unless otherwise noted, all gene and amino acid sequences were retrieved from GenBank (Table S1). Sequence alignments were created using ClustalW and exported using Geneious. Figures were created using assets from www.BioRender.com.

### Peptide synthesis

To generate a C-terminal acid with the last amino acid of the sequence pre-loaded and a density of ~0.2 mmol/g, 150 mg 2-chlorotrityl resin (ChemPep, 100-200 mesh) was washed with DMF and DCM and swelled for 30 min, 0.03 mmol of the respective amino acid was dissolved in 1 mL DMF/DCM and 0.15 mmol DIPEA was added to the amino acid solution. This mixture was added to pre-swelled resin and rotated at RT for 1 hour. Then the resin was washed with 3x DCM and capped with 20 mL 17:2:1 DCM:MeOH:DIPEA with manual mixing between washes. Finally, resin was washed 3x with ~4 mL DCM and 3x with ~4 mL DMF prior to synthesis. Remaining amino acids were chemically synthesized using solid-phase peptide synthesis (SPPS) with a standard synthesis scale of 30 μmol on a Prelude X peptide synthesizer (Gyros Protein Technologies) using Fmoc-protected amino acids (Gyros Protein Technologies). Uroguanylin peptides were synthesized using Fmoc-Cys(Acm)-OH (Aapptec) at C7/C15. The following cycles were used during SPPS: Fmoc deprotection: 3 x 3-min cycles of 2 mL 20% piperidine in DMF. Amino acid coupling: mixing Fmoc-protected amino acid in NMP (0.65 mL, 200 mM), HATU in DMF (0.65 mL, 195 mM), plus NMM in DMF (0.5 mL, 0.6 M) for 25 min with shaking at 150 rpm and nitrogen bubbling. DMF washing (2 mL) was performed between deprotection and coupling steps (3 x 30 s). Completed peptides were washed with DCM and dried prior to cleavage.

### Peptide cleavage from resin

Peptides were cleaved from resin for 2 h using 3.7 mL TFA, 100 μL water, 100 μL TIS and 100 μL EDT. After cleavage, peptides were precipitated into cold diethyl ether (35 mL), stored at −20°C overnight, then washed with diethyl ether (3 x 30 mL), and finally peptides were pelleted by centrifugation and dried overnight in a desiccator prior to HPLC purification.

### Crude peptide purification (STp, STh, Y-STa, V-ST)

Crude peptides were purified by reverse-phase HPLC on a Phenomenex Jupiter 4-μm Proteo C12 90 Å (250 x 21.2 mm) column with a water/ACN gradient in 0.1% TFA (Buffer A: water with 0.1% TFA, Buffer B: 90% ACN with 10% water and 0.1% TFA; for LC/MS buffer is 0.1% formic acid instead of TFA). Specific gradients over 35 min at 10 mL/min were: 29-34% Buffer B (Y-STa and STh), 38-42% Buffer B (V-ST), and 34-38% Buffer B (STp). Purified peptides were lyophilized and their purity validated by analytical HPLC on a Phenomenex Jupiter 4-μm Proteo C12 90 Å (150 x 4.6 mm) column with a linear gradient of 5-90% Buffer B over 30 min. Masses of the peptides were confirmed by LC/MS on an Agilent Technologies 6120 single-quadrupole mass spectrometer with an Agilent 1260 Infinity II HPLC using an Agilent 2.7-μm Poroshell C18 120 Å (4.6 x 50 mm) column.

### Oxidation of free cysteine residues (STp, STh, Y-ST, V-ST)

Oxidation of free cysteine residues in purified peptides was accomplished using air oxidation of solid peptide dissolved in peptide oxidation buffer (50 mM Tris Base, 2% DMSO, pH 8 at 37°C) at ≤0.4 mg/mL (to ensure intramolecular disulfide formation) for ~17 h at 37°C with shaking at 200 rpm. The reaction was quenched with glacial acetic acid (final concentration of 5%), and the pH was confirmed to be between pH 3-4. Reaction solution was then diluted 2-fold with HPLC Buffer A, spun for 10 min at 4696 xg and purified by reverse phase HPLC on a Phenomenex Jupiter 4-μm Proteo C12 column using a gradient of 36-40 % Buffer B (V-ST), 24 to 29% Buffer B (Y-STa), 30-35% Buffer B (STp), 25.5-30% Buffer B (STh) over 35 min with a flow rate of 10 mL/min. All HPLC methods started at 20% Buffer B to prevent peptide precipitation. Fractions were collected and checked for purity by LC/MS (10-90 or 10-70% Buffer B over 10 min) and lyophilized. For each peptide, eight distinct peaks were collected (Figure S5A) and tested for activity in the cellular cGMP assay. The most active peaks were: oxidized STp peak #2, oxidized STh peak #8, oxidized V-ST peak #2, and oxidized Y-ST peak #1 (Figure S5B and C).

### Oxidation of free cysteine residues (human, *P. vampyrus, M. lucifigus*, and *E. fuscus* uroguanylin peptides)

Crude peptides were first validated by LC/MS and analytical HPLC, as described above. Oxidation of free cysteine residues was accomplished using air oxidation of solid peptide dissolved in peptide oxidation buffer at ≤0.4 mg/mL for 14-18 h at 37°C with shaking at 200 rpm. The reaction was quenched with glacial acetic acid (5% final concentration) and the pH was confirmed to be between 3-4. Reaction solution was then diluted 2-fold with HPLC Buffer A, spun for 10 min at 4696 xg and purified by reverse-phase HPLC on a Phenomenex Jupiter 4-μm Proteo C12 column (see specs above) using a gradient of 28-35 % Buffer B (Human), 23-35 % Buffer B (*P. vampyrus*), 22-40% Buffer B (*M. lucifigus*), or 32-50% Bufer B (*E. fuscus*) over 35 min, all HPLC methods starting at 20% Buffer B to prevent peptide precipitation. Fractions were collected and checked for purity by LC/MS (10-90 or 10-70% Buffer B over 10 min, 1 mL/min).

### Acm removal and formation of second disulfide (human, *P. vampyrus, M. lucifigus* uroguanylin peptides)

Dry purified peptides with a single formed disulfide and Cys(Acm) protecting groups at positions C7 and C15 (1-5 mg) were dissolved in Buffer A to a concentration of ~400 μM. Fresh I_2_-TFA mixture was prepared (~10 mg I_2_ dissolved in 5 mL ACN, then added to a solution of 15 mL H_2_O with 0.6 mL TFA), and the I_2_-TFA mix was added (2x volume) to the peptide-Buffer A solution. This mixture reacted on a rotator at RT for 20 min before quenching with 1 M ascorbic acid added drop-wise until solution changes from rust color to colorless, clear solution (typically less than 30 μL). Quenched solution was diluted with one volume Buffer A, spun 10 min at 4696 xg, and transferred to a new tube for purification. Same-day purification proceeded via reverse-phase HPLC using a Phenomenex 5-μm C18-Kinetex column C10 100Å (150 x 4.6 mm) with a gradient of 20-52% Buffer B (human uroguanylin), 14-23% Buffer B (*P. vampyrus*), 10-30% Buffer B (*M. lucifugus*), or 22-30% Buffer B (*E. fuscus*) over 30 min at 2 mL/min to achieve baseline separation of peptide topoisomers. Two individual peaks were collected separately and their mass confirmed by LC/MS. Generally the larger peak was determined to be the active isomer (consistent with previous studies^24^). The uroguanylin topoisomers were found to interconvert at RT (as seen previously for human uroguanylin^24–26^), with *M. lucifigus* converting the fastest—within 30 min of peak purification). Therefore, fractions were placed on ice immediately after purification until confirmation by LC/MS, then pooled and lyophilized. All dry, purified peptides were stored in parafilmed containers in the dark at −20°C. We tested the order of disulfide bond formation and found that forming the C7-C15 disulfide second yielded more of the active isomer compared to forming the C4-C12 disulfide second.

### Construction of transgenic 293T cell lines

To generate each GC-C expressing cell line 2.5 x 10^5^ 293T cells were seeded into a single well of a 6 well plate. After 24 hours of growth, 50 μL concentrated lentivirus encoding GC-C linked to a GFP reporter was added to a total of 1 mL growth medium containing 8 μM polybrene (Sigma). 24 hours post transduction, cells were transferred to a T75 flask and grown for 5-7 days. Transgenic lines were established by FACS sorting the top 25% of GFP expressing cells from each transduction.

### Cellular cGMP assay

293T cells expressing GC-C variants were grown to confluence in 24-well plates before ligand stimulation. Toxin and uroguanylin solutions were diluted in reduced serum media (Opti-MEM, Thermo Scientific) to the appropriate experimental concentration. For each measurement, cell culture media was then aspirated and replaced with the ligand-containing solution in 3-6 replicate wells. Following incubation at 37°C/5% CO_2_ for 20 min, the ligand solution was aspirated and cells lysed directly in 0.1 M HCl. Intracellular cGMP was then measured using an Enzyme Linked Immunosorbent Assay kit (Enzo Scientific) according to the manufacturer’s specifications. Absorbance was measured using a plate reader (BioTek), and cGMP concentrations calculated based on a standard curve. Four-parameter variable slope dose-response curves were generated in Prism 8 (GraphPad).

### Lentivirus generation and cloning

GC-C sequences were downloaded from GenBank based on whole genome sequences from each species tested. Each C-terminally HA tagged GC-C variant was synthesized via gene synthesis (Life Technologies). GC-C was then cloned into the lentiviral transfer vector pUltra (Addgene #24129) between the XbaI and BamHI restriction sites by Gibson Assembly, in frame with GFP and the T2A linker sequence. To generate lentiviral particles, 10 cm dishes were seeded with 3 x 10^6^ 293T cells 24 hours prior to transfection. Cells were then transfected with 7.6 μg pUltra-GC-C, 7.6 μg psPAX2 packaging plasmid (Addgene # 12260), and 3.8 μg pMD2.G envelope plasmid (Addgene # 12259) with 56 μl FuGene HD transfection reagent according to the manufacturer’s specifications. Media was replaced 24 hours post-transfection and replaced with 10 mL media. Viral supernatants were collected 48 hours-post transfection and passed through a 0.4 μm followed by overnight incubation with 1X PEG-IT solution (System Biosciences) at 4°C. Precipitated viral particles were centrifuged at 1500xg for 30 min at 4°C and resuspended in PBS at a final volume of 500 μl before storage at −80°C.

### Western blotting

293T cells expressing GC-C were grown to confluence in 24-well plates following immunoblot analysis. To avoid aggregation, cells were lysed directly in 2x Laemmli buffer containing 8 M urea (Sigma), 3 M thiourea (Sigma), and β-mercaptoethanol for 30 min at room temperature prior to analysis. Total protein was resolved by Mini-PROTEAN GTX polyacrylamide gel electrophoresis (Bio-rad). Proteins were detected using anti-HA (Covance cat# MMS-101P, 1:1000) and anti-Actin (BD cat# 612657, 1:1000) antibodies. Blots were visualized using film or C-DiGit chemiluminescent imager (LI-COR).

### Intestinal organoid culture

Small intestinal organoids derived from GC-C knockout mice were a generous gift from the laboratory of Scott Waldman and were isolated as described previously ^21^. Organoids were maintained in 50 μl Matrigel droplets (Corning) in wells of a 24-well plate containing 650 μl IntestiCult mouse organoid growth medium (StemCell Technologies cat# 06005) at 37°C/5% CO_2_. Organoids were passaged every 7-10 days by disruption with TrypLE (Thermo Scientific) and reseeded at a concentration of 100-500 organoids per well.

### Intestinal organoid transduction

For each transduction approximately 500 organoids were mechanically disrupted by vigorous pipetting using a 20 μl pipette tip. Disrupted organoids were then resuspended in 100 μl concentrated lentivirus solution containing 8 μM polybrene (Sigma), 10 μM Y-27632 (Sigma) and 10 μM SB431542 (Sigma). Lentiviral infection was allowed to proceed at 37°C/5% CO_2_ for 6 h prior to resuspension in Matrigel and seeding into a new well. Transduced organoids were monitored for GFP expression and 20-50 organoids with uniform fluorescence were selected manually 72 h post-transduction and transferred to a new well to establish transgenic lines.

### Organoid swelling assay

Organoids were grown in a 24 well plate for 5-6 days after passaging prior to swelling assays at a concentration of 100-500 organoids per well. Organoids were imaged on an ImageXPress Pico Automated Cell Imaging System (Molecular Devices) using the live cell imaging cassette with temperature set to 37°C, 5% CO_2_, and humidity levels 85-95%. These parameters were monitored and remained constant throughout each experiment. For swelling analysis, ST toxins were added directly to the organoid culture media to a final concentration of 5 μM. Following addition of toxin, a 24 mm^2^ area of each well (corresponding to ~13% of the total well area) was imaged every 10 min at 4x magnification for the full 2 hours of the experiment. The imaged area was constant across the experiment and analogous between wells and contained at least 50 organoids for each analyzed strain. 20 stacked transmitted light images were collected for each well with a focus step of 25 μm and the Best Plane image was calculated by the ImageXPress software. GFP images were collected at the beginning and end of the experiment using the same imaging parameters, and both the best focus and maximum fluorescence images for each stack were calculated and output by the ImageXPress software.

Organoids used for analysis were identified in ImageJ and assigned ROI identifiers so that the same organoids could be easily assessed across timepoints. 30 organoids from each strain were selected randomly to include a variety of starting sizes and were checked for GC-C expression by comparison with GFP images. GC-C expressing organoids that ruptured during the experiment were excluded from analysis. Swelling was assessed by measuring the area of each organoid in ImageJ at t=0 and t=120 min. The percent change in organoid area from 0 to 120 min was calculated for each organoid and plotted in Prism. Significance in organoid swelling data was assessed by an unpaired t-test with Welch’s correction using Prism 8.

---

**Table S1:**
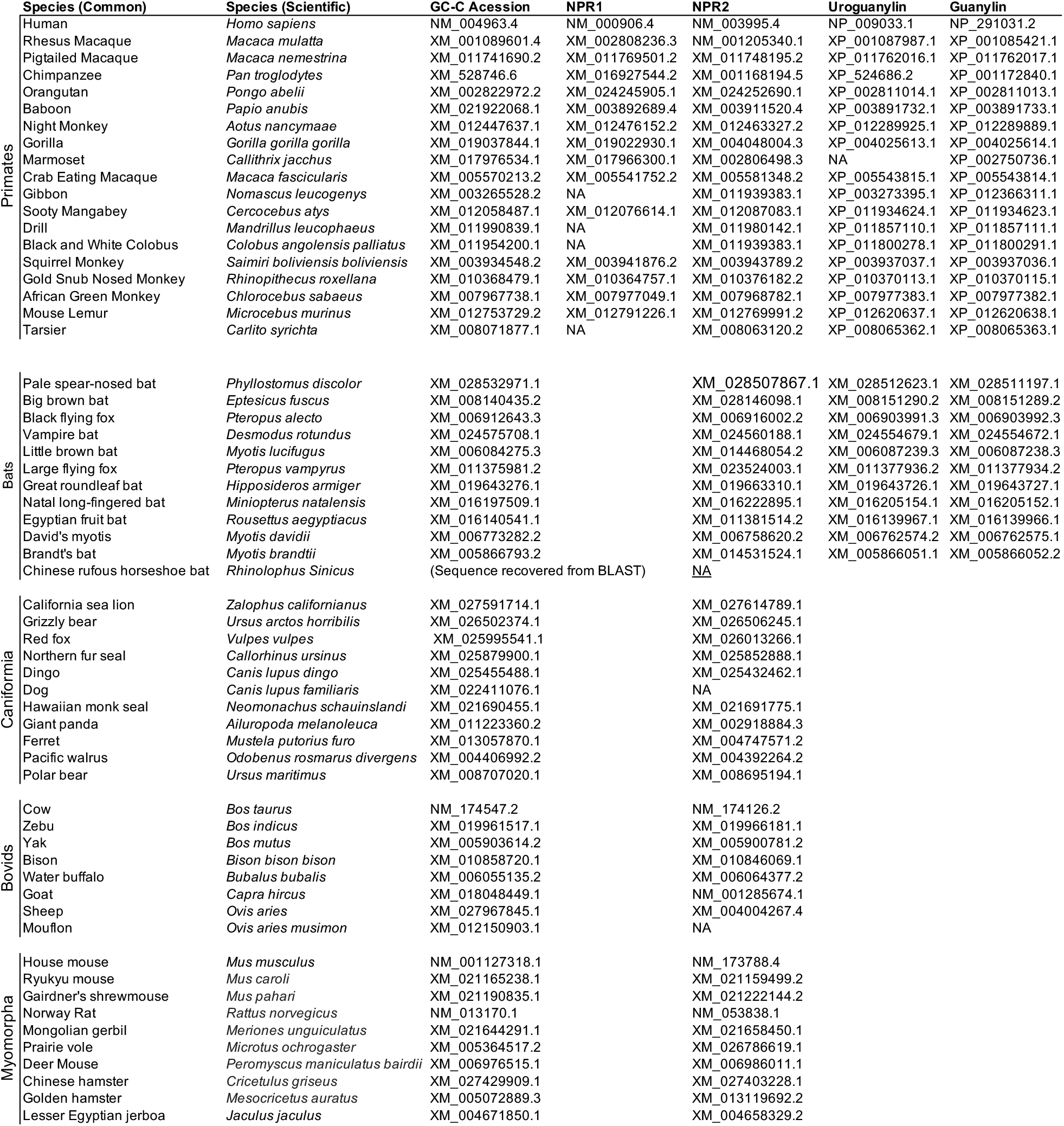
Gene sequences used in this study

**Table S2:**
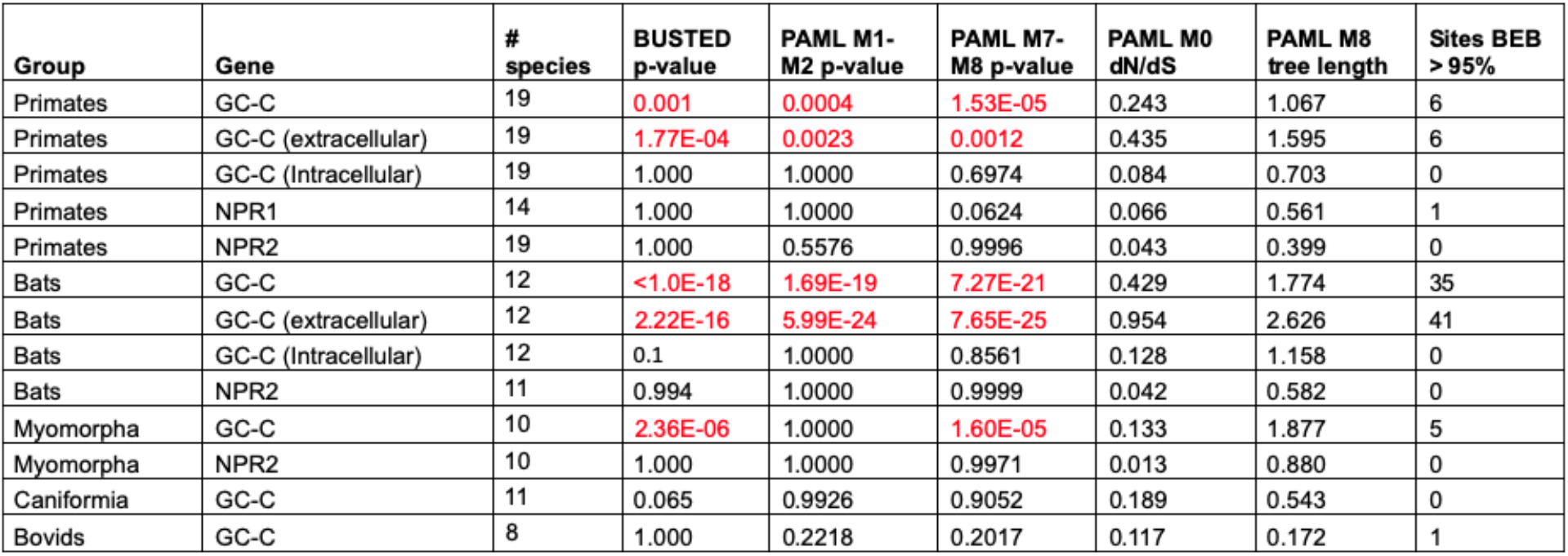
Likelihood ratio tests for positive selection in mammalian GC-C and NPR genes.

**Figure S1:**
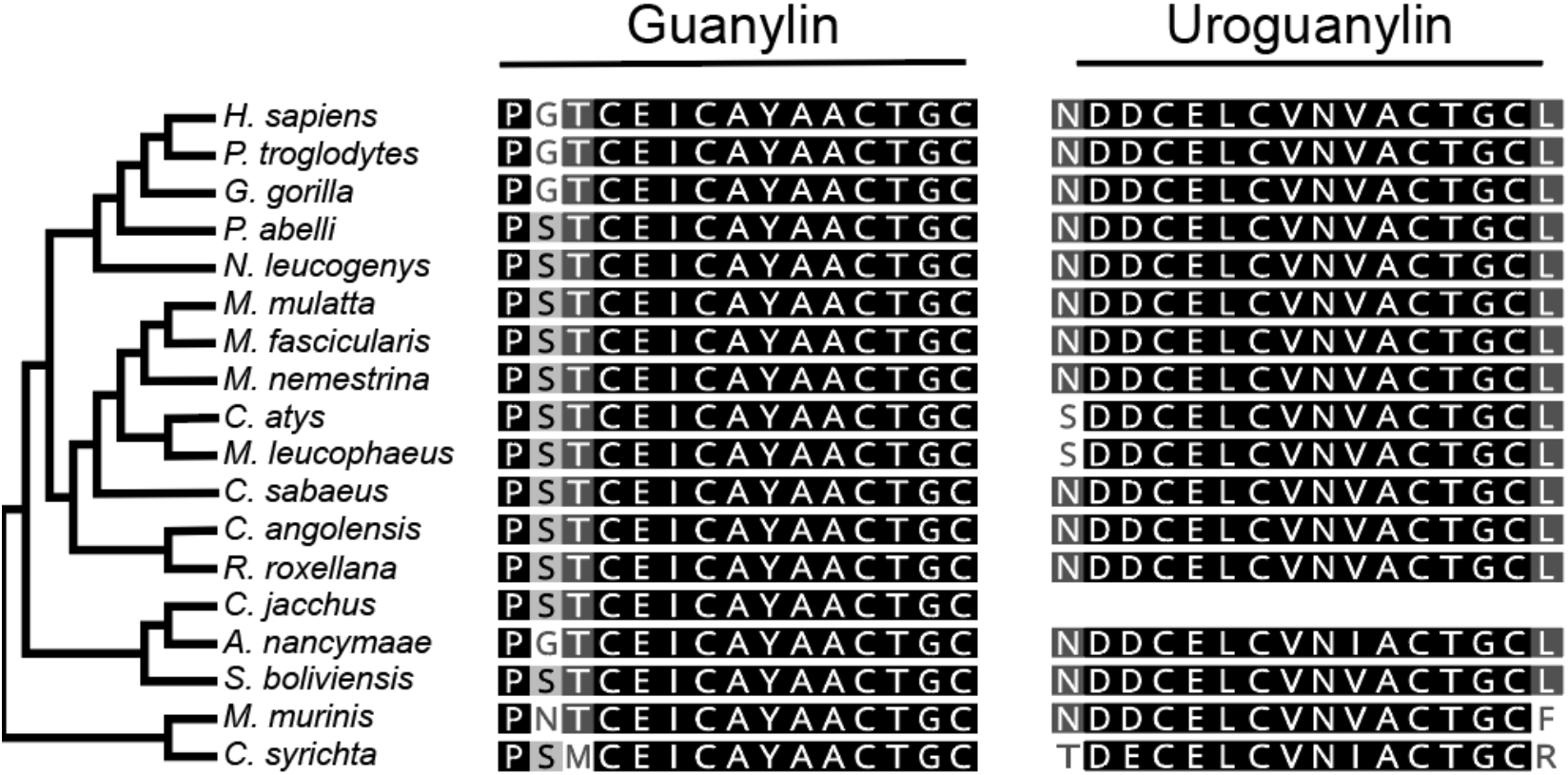
Sequence conservation in primate guanylin and uroguanylin. Amino acid sequence alignments of the secreted forms of primate guanylin and uroguanylin are shown. Sequences were retrieved from genbank and aligned using clustal omega. The sequence of marmoset uroguanylin (*C. jacchus*), was unavailable.

**Figure S2:**
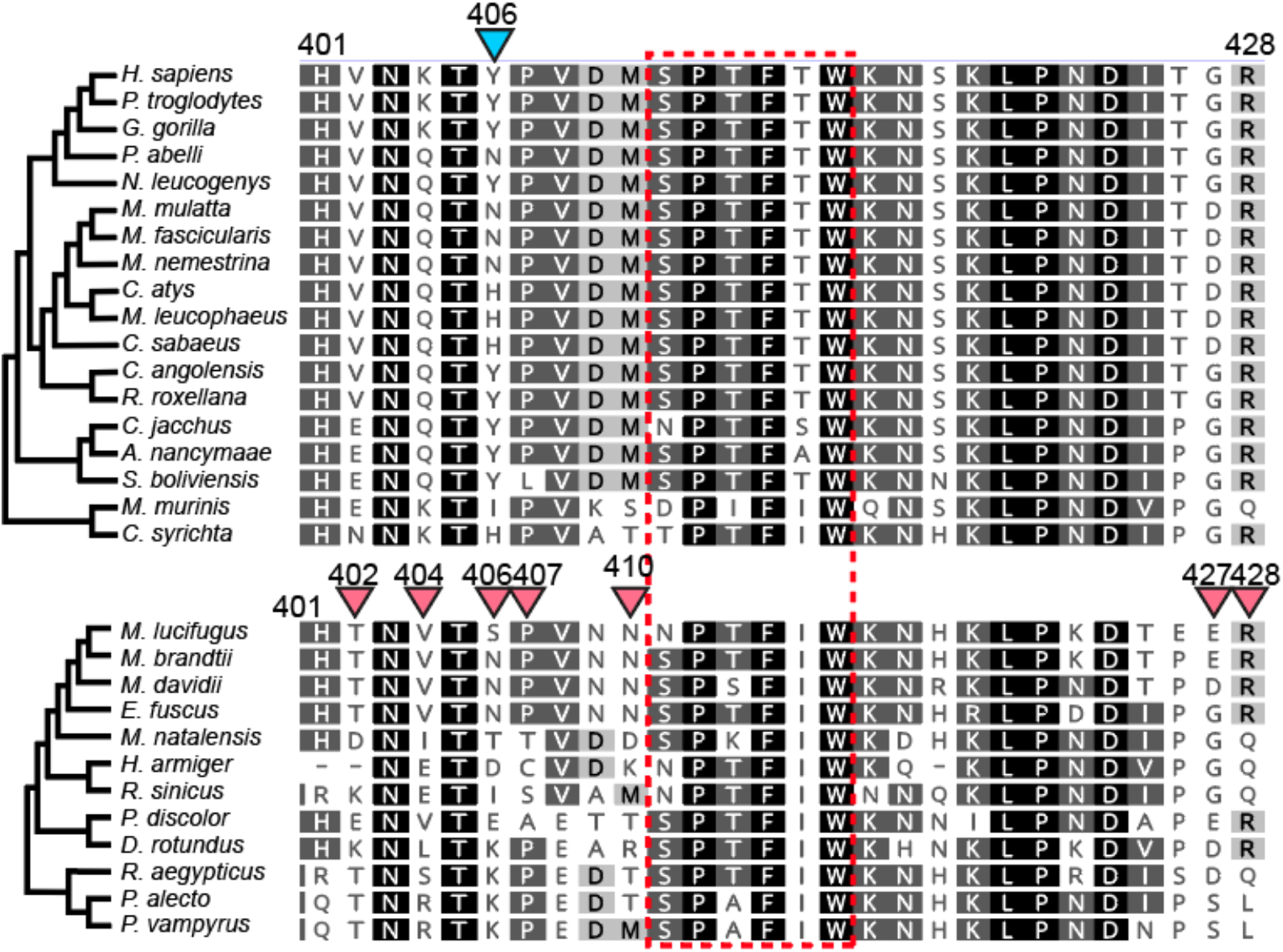
Positive selection detected near the putative GC-C STa-binding pocket. GC-C amino acid alignments are shown from primates (top) and bats (bottom) from amino acid 400-428. A proposed ligand-binding pocket is highlighted^13^ (red dashed box). Amino acid positions with significantly elevated dN/dS ratios are highlighted with blue (primates) or red (bats) triangles (PAML >95% posterior probability).

**Figure S3:**
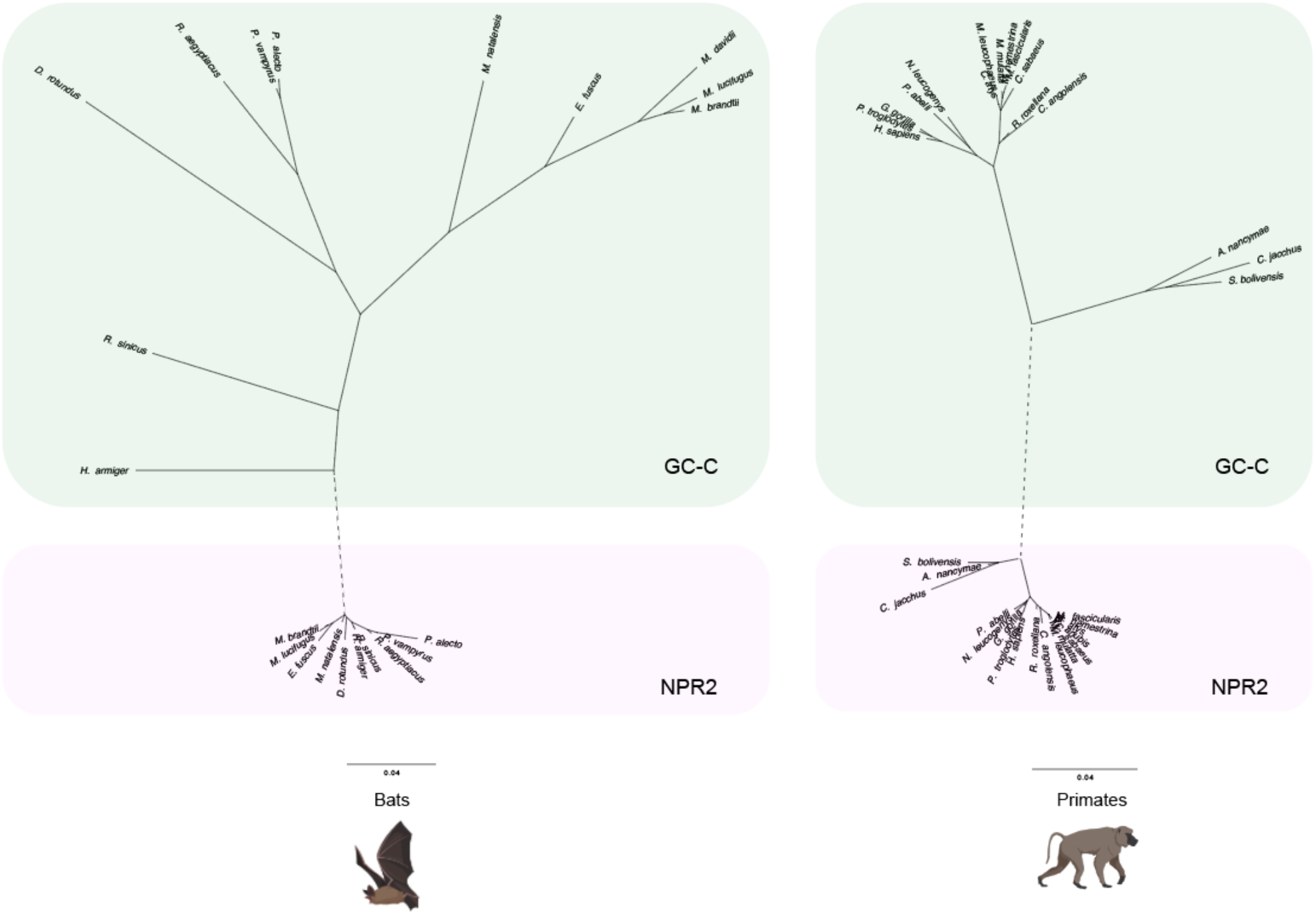
GC-C ligand-binding domain sequence divergence in primates and bats. Amino acid alignments were generated for the ligand-binding domains of GC-C and NPR2 in primates and bats. Trees were generated using PHYML with the Le Gascuel amino acid substitution model. Branches connecting NPR2 and GC-C trees are artificially collapsed (dashed line).

**Table S3:** Positively selected amino acid positions in primate and bat GC-C identified by PAML

|  | Sites with dN/dS > 1 * | Probability |
| --- | --- | --- |
| Primate GC-C | S21 | 0.909 |
|  | F22 | **0.966** |
|  | S23 | 0.908 |
|  | Q25 | **0.975** |
|  | V224 | **0.983** |
|  | G251 | 0.939 |
|  | Q279 | **0.965** |
|  | R312 | 0.923 |
|  | T317 | 0.93 |
|  | D364 | **0.99** |
|  | W372 | 0.927 |
|  | Y406 | **0.985** |
|  | S419 | 0.928 |
|  | R688 | 0.915 |
| Bat GC-C | I 21 | **0.99** |
|  | W 22 | **0.981** |
|  | T 23 | **0.997** |
|  | S 24 | **0.996** |
|  | P 25 | **0.977** |
|  | S 27 | **0.997** |
|  | H 31 | **0.975** |
|  | I 34 | **0.971** |
|  | Q 48 | **0.995** |
|  | E 52 | **0.997** |
|  | E 60 | **0.966** |
|  | R 66 | **0.964** |
|  | E 67 | **0.996** |
|  | I 70 | **0.989** |
|  | N 71 | 0.917 |
|  | S 72 | 0.901 |
|  | S 79 | 0.901 |
|  | S 81 | **0.955** |
|  | Y 84 | 0.909 |
|  | P 88 | **0.998** |
|  | V 93 | 0.929 |
|  | T 111 | **0.98** |
|  | K 114 | **0.986** |
|  | A 177 | **0.99** |
|  | S 186 | 0.913 |
|  | N219 | 0.941 |
|  | N 221 | **0.992** |
|  | K 230 | 0.944 |
|  | H 239 | 0.913 |
|  | N 240 | 0.944 |
|  | T 251 | **0.989** |
|  | T 254 | 0.937 |
|  | S 256 | **0.998** |
|  | I 257 | 0.914 |
|  | Q 282 | 0.929 |
|  | S 300 | 0.927 |
|  | T 304 | **0.972** |
|  | I 308 | **0.977** |
|  | S 309 | 0.915 |
|  | M 318 | **0.951** |
|  | D 345 | 0.933 |
|  | L 349 | **0.989** |
|  | E 402 | **0.992** |
|  | V 404 | **0.996** |
|  | E 406 | **0.985** |
|  | A 407 | **0.952** |
|  | T 410 | **0.99** |
|  | N 419 | 0.942 |
|  | N 423 | 0.932 |
|  | A 425 | 0.937 |
|  | E 427 | **0.993** |
|  | R 428 | **0.958** |
|  | N 744 | 0.944 |
|  | R 993 | 0.94 |

\*Posterior probabilities >90% by PAML Model 8 Bayes Empirical Bayes (BEB) analysis, probabilities >95% are bolded amino acid identities are shown for *H. sapiens* and *M. lucifugus* in primates and bats, respectively

**Figure S4:**
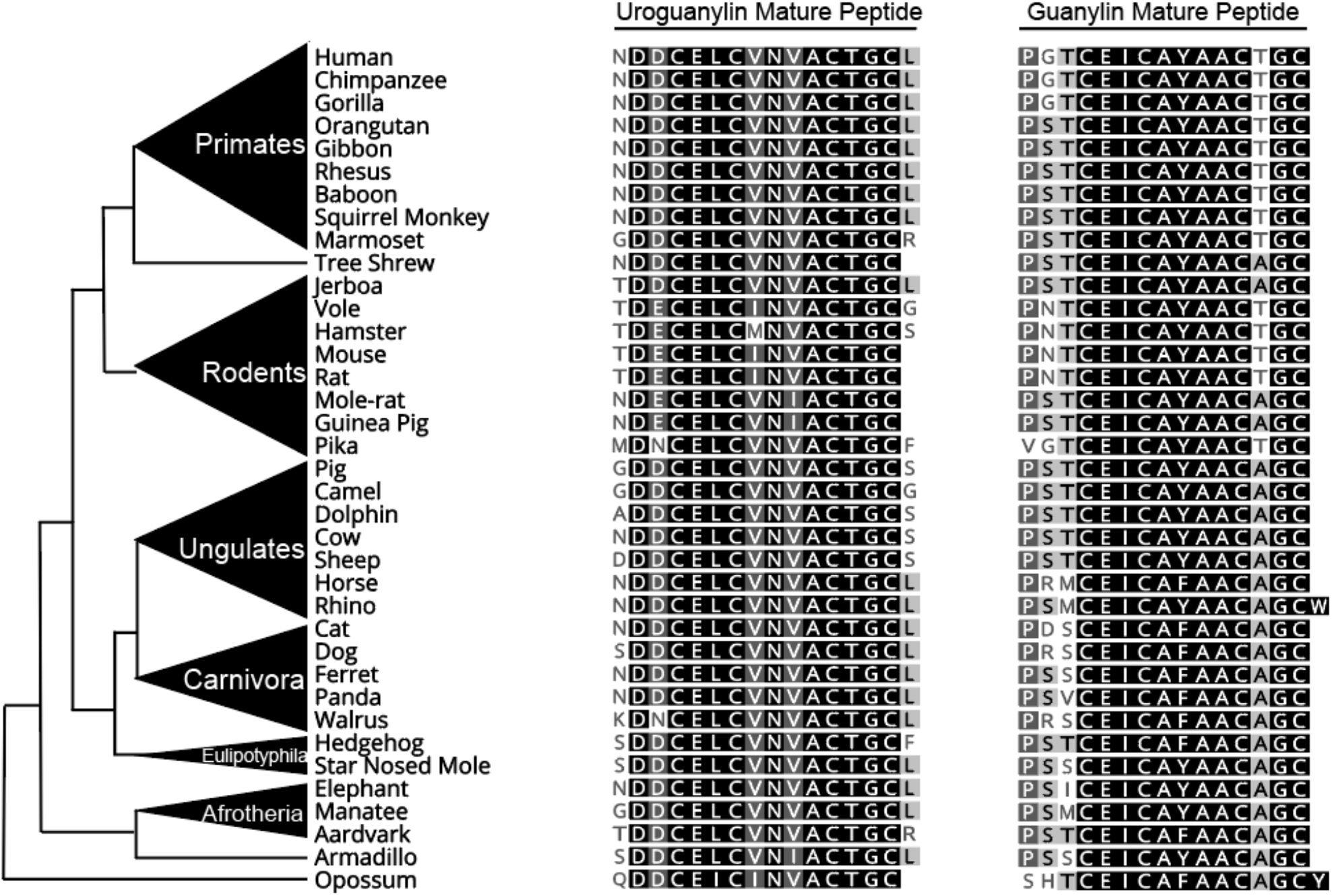
Sequence divergence in endogenous guanylin and uroguanylin peptides in mammals.

**Figure S5:**
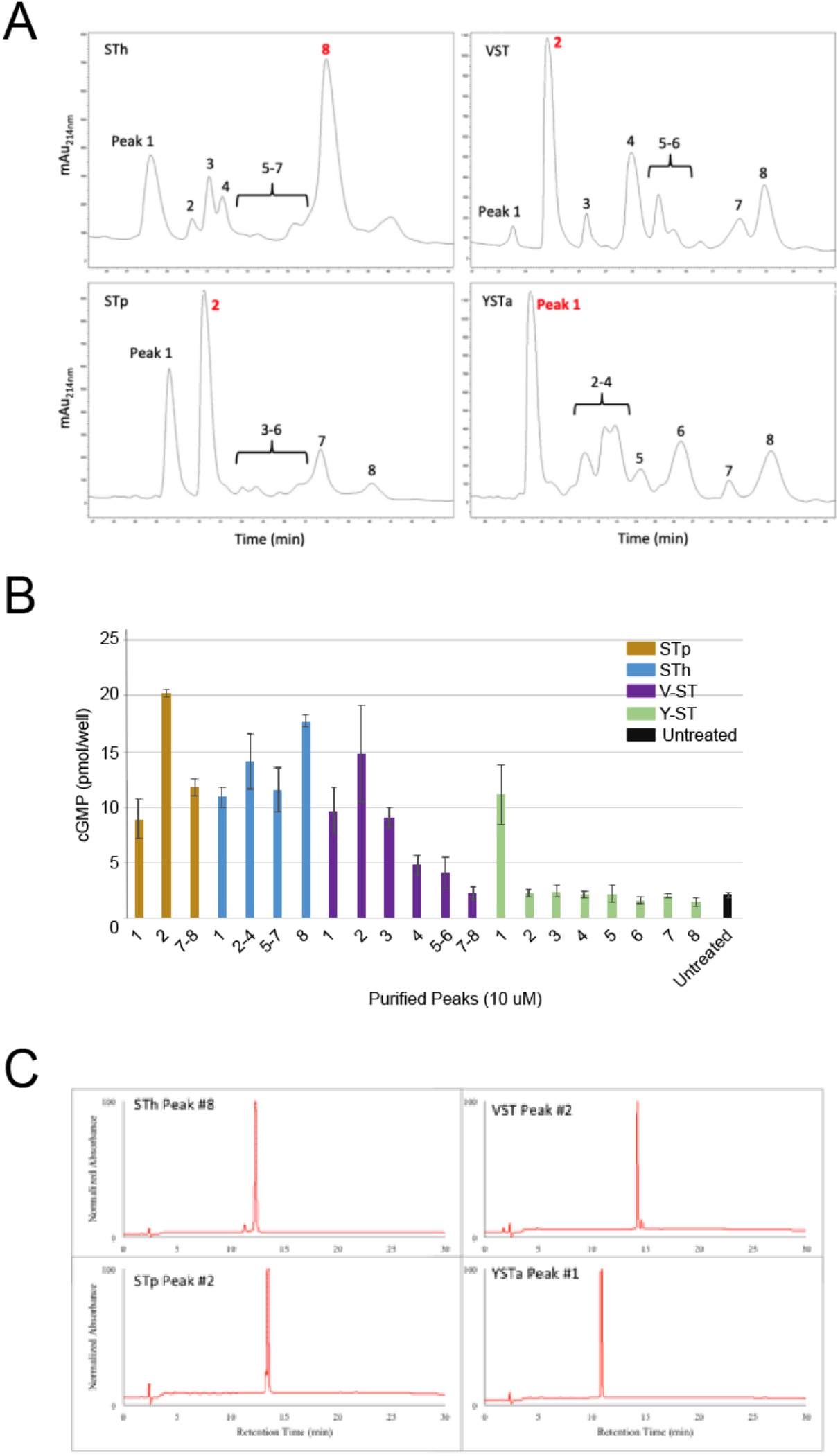
Synthesis and oxidation of active STa peptides. (A) HPLC traces of each synthesized peptide shows evidence of ~8 peaks following air oxidation. Each peak was then collected for downstream analysis. (B) HEK293T cells expressing human GC-C were stimulated with the major purification products to identify the most potent fractions by measuring intracellular cGMP. For each peptide, the predominant oxidation product represented the most potent GC-C ligand. Some peaks were unable to be purified in isolation. (C) Final HPLC purification of toxin peptides used for analysis.

**Figure S6:**
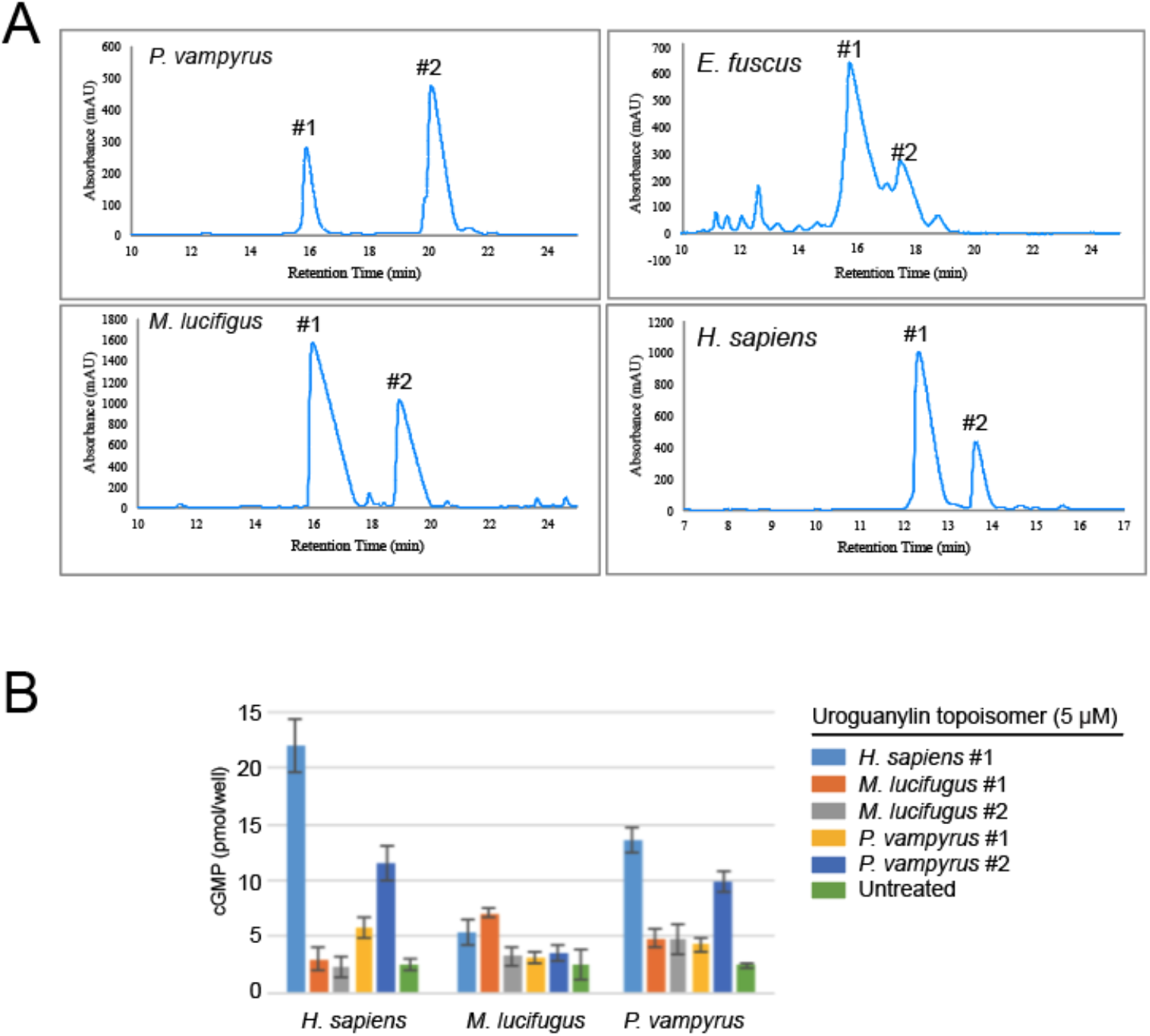
Synthesis and purification of uroguanylin topoisomers. (A)HPLC traces of synthesized uroguanylin from the indicated species. Two distinct topoisomers were observed following oxidation of synthesized uroguanylin from each species. (B) cGMP generation assays in HEK293T cells expressing GC-C from the indicated species were performed to identify the most active topoisomer. Human isomer #1 is the known active peak. *E. fuscus* peak #1 was used in in experimental assays (Figure 4).

